# Alpha-synuclein antisense oligonucleotides as a disease-modifying therapy for Parkinson’s disease

**DOI:** 10.1101/830554

**Authors:** Tracy A. Cole, Hien Zhao, Timothy J. Collier, Ivette Sandoval, Caryl E. Sortwell, Kathy Steece-Collier, Brian F. Daley, Alix Booms, Jack Lipton, Mackenzie Welch, Melissa Berman, Luke Jandreski, Danielle Graham, Andreas Weihofen, Stephanie Celano, Emily Schulz, Allyson Cole-Strauss, Esteban Luna, Duc Quach, Apoorva Mohan, C. Frank Bennett, Eric E. Swayze, Holly B. Kordasiewicz, Kelvin C. Luk, Katrina L. Paumier

## Abstract

Parkinson’s disease (PD) is a prevalent neurodegenerative disease with no approved disease-modifying therapies. Multiplications, mutations, and single nucleotide polymorphisms in the *SNCA* gene, encoding alpha-synuclein protein (aSyn), either cause or increase risk for PD. Intracellular accumulations of aSyn are pathological hallmarks of PD. Taken together, reduction of aSyn production may provide a disease-modifying therapy for PD. We show that antisense oligonucleotides (ASOs) reduce production of aSyn in rodent pre-formed fibril (PFF) models of PD. Reduced aSyn production leads to prevention and removal of established aSyn pathology and prevents dopaminergic cell dysfunction. In addition, we address the translational potential of the approach through characterization of human *SNCA* targeting ASOs that efficiently suppress the human *SNCA* transcript *in vivo.* We demonstrate broad activity and distribution of the human *SNCA* ASOs throughout the non-human primate brain and a corresponding decrease in aSyn cerebral spinal fluid (CSF) levels. Taken together, these data suggest that by inhibiting production of aSyn it may be possible to reverse established pathology and thus supports the development of *SNCA* ASOs as a potentially disease modifying therapy for PD and related synucleinopathies.

**Summary:** Antisense oligonucleotides designed against *SNCA*, which are progressing to the clinic, have the potential to be a disease modifying therapeutic for Parkinson’s disease patients.

## Introduction

There is strong genetic evidence implicating the role of aSyn (alpha-synuclein protein) in the pathogenesis of Parkinson’s disease (PD) (Chartier-Harlin et al., 2004; Devine et al., 2011; Farrer et al., 2004; Fuchs et al., 2007; Fuchs et al., 2008; Ibanez et al., 2009; Kruger et al., 1998; Mata et al., 2010; Mutez et al., 2011; Ross et al., 2008; Satake et al., 2009; Simon-Sanchez et al., 2009; Singleton et al., 2003; Soldner et al., 2016; Spillantini et al., 1997). Intracellular aSyn aggregates are a pathological hallmark of PD that increase in number and spread through the brain as symptoms worsen in PD patients (Goedert, 2015; Goedert et al., 2013). Duplication, triplication or genetic mutations in the *SNCA* gene (produces aSyn protein) (A53T, A30P, E46K, G51D, etc.) are linked to autosomal dominant forms of the disease (Chartier-Harlin et al., 2004; Farrer et al., 2004; Fuchs et al., 2007; Miller et al., 2004; Ross et al., 2008; Singleton et al., 2003). Moreover, polymorphisms that occur within specific regions of the *SNCA* gene increase the overall risk of PD by either increasing the production or slowing the clearance of aSyn (Cronin et al., 2009; Fuchs et al., 2008; Mata et al., 2010; Nalls et al., 2014; Simon-Sanchez et al., 2009; Soldner et al., 2016). A toxic gain of function of aSyn is also established in other synucleinopathies including multiple systems atrophy (MSA) (Spillantini et al., 1998), Diffuse Lewy body disease (DLBD) (Nishioka et al., 2010), and Gaucher disease (GD) (Aflaki et al., 2017), which collectively affects about 1% of people over 60 years of age. Clinically diagnosed Dementia with Lewy bodies (DLB) (Spillantini et al., 1998) and pure autonomic failure (PAF) (Arai et al., 2000; Isonaka et al., 2017) also exhibit Lewy pathology, suggesting a toxic gain of function of aSyn in DLB and PAF.

To date, there are multiple therapeutic strategies being investigated, including antibodies and small molecule approaches targeting different forms and conformational states of aSyn, however, the toxic species of aSyn has not yet been confirmed, potentially limiting therapeutic benefit of these approaches (Dehay et al., 2015; Kingwell, 2017). In addition, antibodies and small molecules often only target extracellular pools of the protein. However, antisense oligonucleotide (ASO) therapy can potentially overcome the limitations of these approaches since they inhibit the production of aSyn by targeting RNA intracellularly, thereby reducing all forms of the aSyn protein (Bennett and Swayze, 2010).

To determine the efficacy of ASOs in synucleiopathies, we use rat and mouse aSyn pre-formed fibril (PFF) transmission models, which replicate aspects of human PD progression including seeding and aggregate deposition of endogenous aSyn, reduced striatal dopamine, dopaminergic cell dysfunction (tyrosine hydroxylase (TH) loss) in the substantia nigra, and motor dysfunction (Luk et al., 2012; Paumier et al., 2015). We hypothesize that these pathological changes can be prevented or reversed by inhibiting the production of aSyn using ASOs targeted to the *Snca* gene. Though aSyn pathology has been shown to be reversible with complete genetic ablation in adult mice in overexpression and toxin models (Hayashita-Kinoh et al., 2006; Lim et al., 2011; Masliah et al., 2005; Tran et al., 2014; Uehara et al., 2019; Zharikov et al., 2015) and with suppression in cells (Luna et al., 2018), we aimed to determine if partial transient and/or sustained suppression of endogenous aSyn with a therapeutically relevant approach *in vivo* could lead to reversal of established pathology and prevention of TH loss. We also aimed to improve cellular function, of which some aspects have been shown to be dysfunctional in iPSC-derived PD patient neurons, PD patient tissue, as well as in animal models of PD (Di Maio et al., 2016; Ludtmann et al., 2018; Martin et al., 2006; Muller et al., 2013; Tapias et al., 2017). Remarkably, we found a robust clearance of aSyn pathology when production of aSyn is inhibited, which results in improvement in cell function as measured by double strand DNA breaks. With human *SNCA* targeting ASOs, we demonstrate widespread target engagement in the brain of transgenic mice expressing human *SNCA* and in all of the PD relevant brain regions in NHPs, thus demonstrating the therapeutic potential of ASOs for PD and other synucleinopathies.

## Results

### Reduction of *Snca* improves cellular function in cells and prevents pathogenic aSyn aggregate deposition in an *in vivo* PFF model of PD

Two 2’-O-methoxyethyl/DNA gapmer ASOs targeting *Snca* were used for *in vitro* and *in vivo* experimentation in addition to a control ASO (Table 1A). In previous work, ASO suppression of *Snca* in a primary cortical culture PFF system inhibited *Snca* mRNA and reversed pSer129+ pathology and cellular dysfunction (Luna et al., 2018). To examine the capability of ASOs to improve cellular function we utilized this same ASO (ASO1) and system. As cellular function is impaired in PD we sought to characterize DNA double strand breaks, an increase in which indicates dysfunctional DNA repair, to determine whether a *Snca* ASO could mitigate dysfunction. As expected, application of PFFs resulted in production of pSer129+ aggregates in primary mouse cortical cultures and this led to double strand breaks (γH2AX Ser139) (Fig. 1A and B). Remarkably, pSer129+ pathology was prevented and double strand breaks (γH2AX Ser139) were normalized to control levels with application of *Snca* ASO1, but not a control ASO (Fig. 1A and B).

**Fig. 1.**
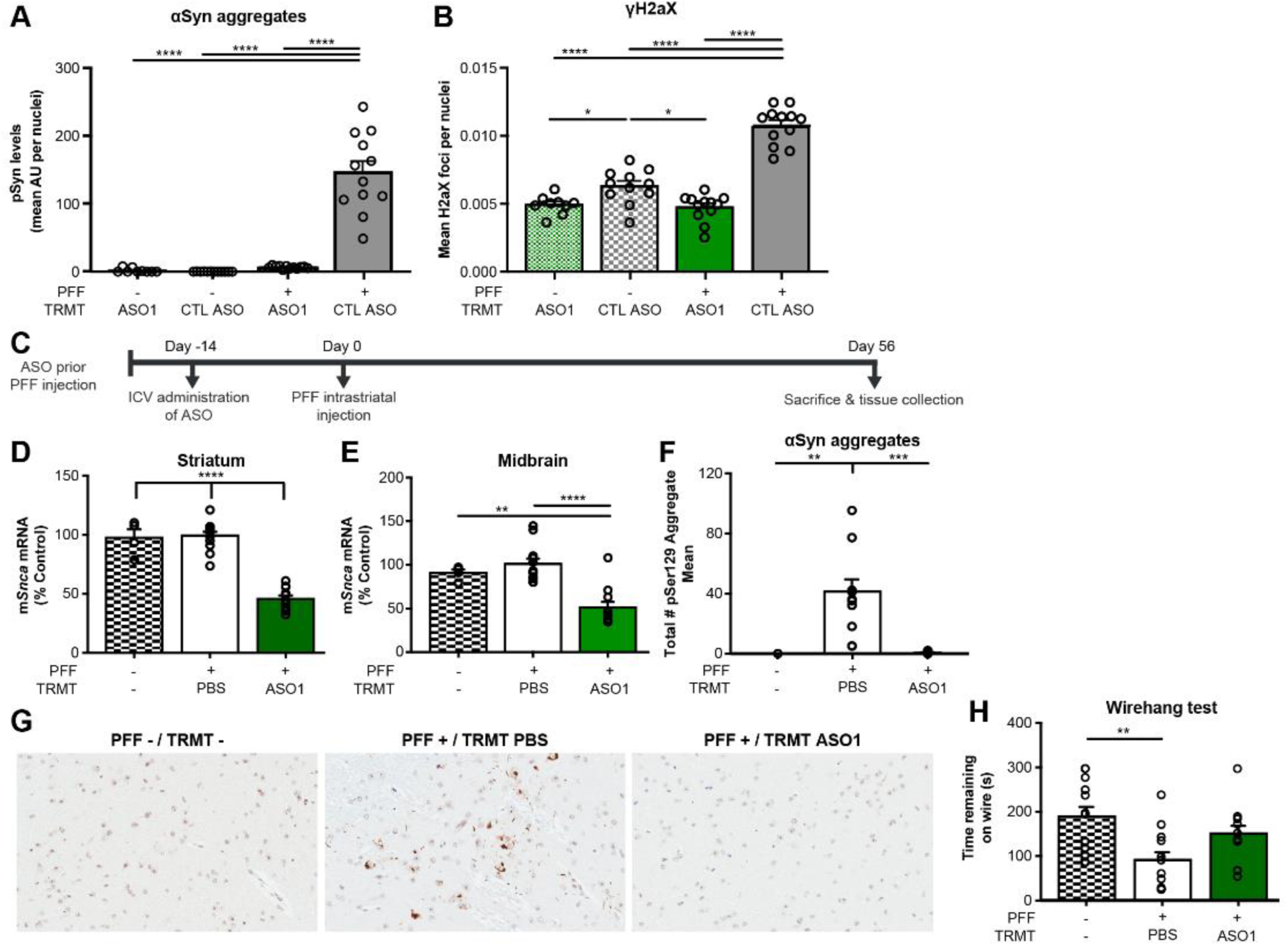
ASO-mediated reduction of *Snca* improves cellular function in cells and prevents pathogenic aSyn aggregate deposition in an *in vivo* PFF model of PD. (**A**) Quantification of pSer129+ area by intensity in mouse primary cortical cultures and (**B**) cellular function, by γH2AX Ser139, in mouse primary cortical cultures with either PBS, CTL ASO, or ASO1 30 minutes following PFF addition. Replicated 2 times. (**C**) Timeline for single 700µg ICV bolus ASO administration prior to PFF injection paradigm with termination at day 56. (**D** to **E**) mRNA reduction by RT-PCR in striatum and midbrain. (**F**) Quantification of pSyn+ aggregate reduction (total enumeration) in the substantia nigra by IHC. (**G**) Representative images of immunostaining (IHC) for pSer129+ aggregate counts. (**H**) Performance on a wire hang task (n=4, 12 and 11 for naïve, PBS and ASO1, respectively, except wire hang, in which n=12 for naive). Error bars represent ± s.e.m. *P<0.05, **P<0.01, ***P<0.001, ****P<0.0001 (Two-way ANOVA with Tukey post hoc analyses for duration of action with all other analyses using One-way ANOVA with Tukey post hoc analyses). PFF (pre-formed fibril), TRMT(treatment).

**Table 1.**
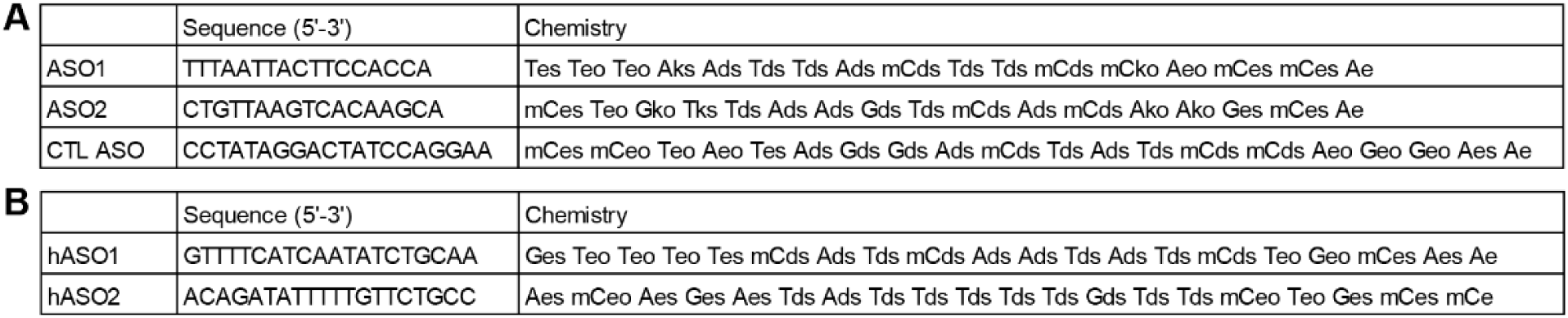
Table of *Snca/SNCA* ASOs used for *in vivo* testing.

To further extend this work, we assessed *Snca* ASOs in a rodent model of PD. Rodent aSyn PFF intrastriatal injection models result in accumulation of phospho-S129 (pSer129+) aggregates that propagate to interconnected regions (including from the substantia nigra (SN) to the striatum) leading to dysfunction of the nigrostriatal system (Luk et al., 2012; Paumier et al., 2015). A single 700µg ICV injection of *Snca*-targeted ASO1 reduced *Snca* mRNA by ∼50% 70 days after ASO administration (Fig. 1, C to E), this resulted in prevention of pSer129+ aggregate deposition in the PFF mouse model (∼96% reduction, Fig 1F and G) 56 days following PFF injection. PFF injected mice exhibit significant deficits in a wire hang motor function task, as published previously (Luk et al., 2012). Following administration of ASO1, mice that were injected with PFFs no longer exhibit a significant deficit in comparison to naïve mice (Fig. 1H).

### ASO-mediated reduction of *Snca* is dose-responsive, exhibits a prolonged duration of action, and dose-responsively prevents pathogenic aSyn aggregate deposition in an *in vivo* PFF model of PD

More extensive characterization of ASO1 demonstrated dose-dependent reduction of *Snca* mRNA *in vivo* 21 days following a single intracerebroventricular (ICV) administration in rats (Fig. 2, A to C). Dose-dependent ASO-mediated *Snca* mRNA suppression with ASO1 also resulted in dose-dependent prevention of pSer129+ aggregate deposition in comparison to the PBS group at 61 days post PFF injection (82 days post ASO ICV administration) in the SN (Fig. 2, D to F and fig. S1, A and B). This is consistent with complete prevention of aggregates in aSyn KO mice and attenuated aggregation in heterozygous KO mice injected with PFFs (Luk et al., 2012). There was no evidence of dopaminergic cell dysfunction at this time point (fig. S1C) as expected (Paumier et al., 2015) and the control ASO exhibited a profile similar to PBS administered rats, indicating the control ASO does not alter disease (Fig. 2, E to F and fig. S1, A to C). ASO1 reduced *Snca* mRNA and prevented pSer129+ aggregate deposition in the SN and across multiple brain regions, including prefrontal cortex and motor cortex 61 days post PFF injection (fig. S1, D to G).

**Fig. 2.**
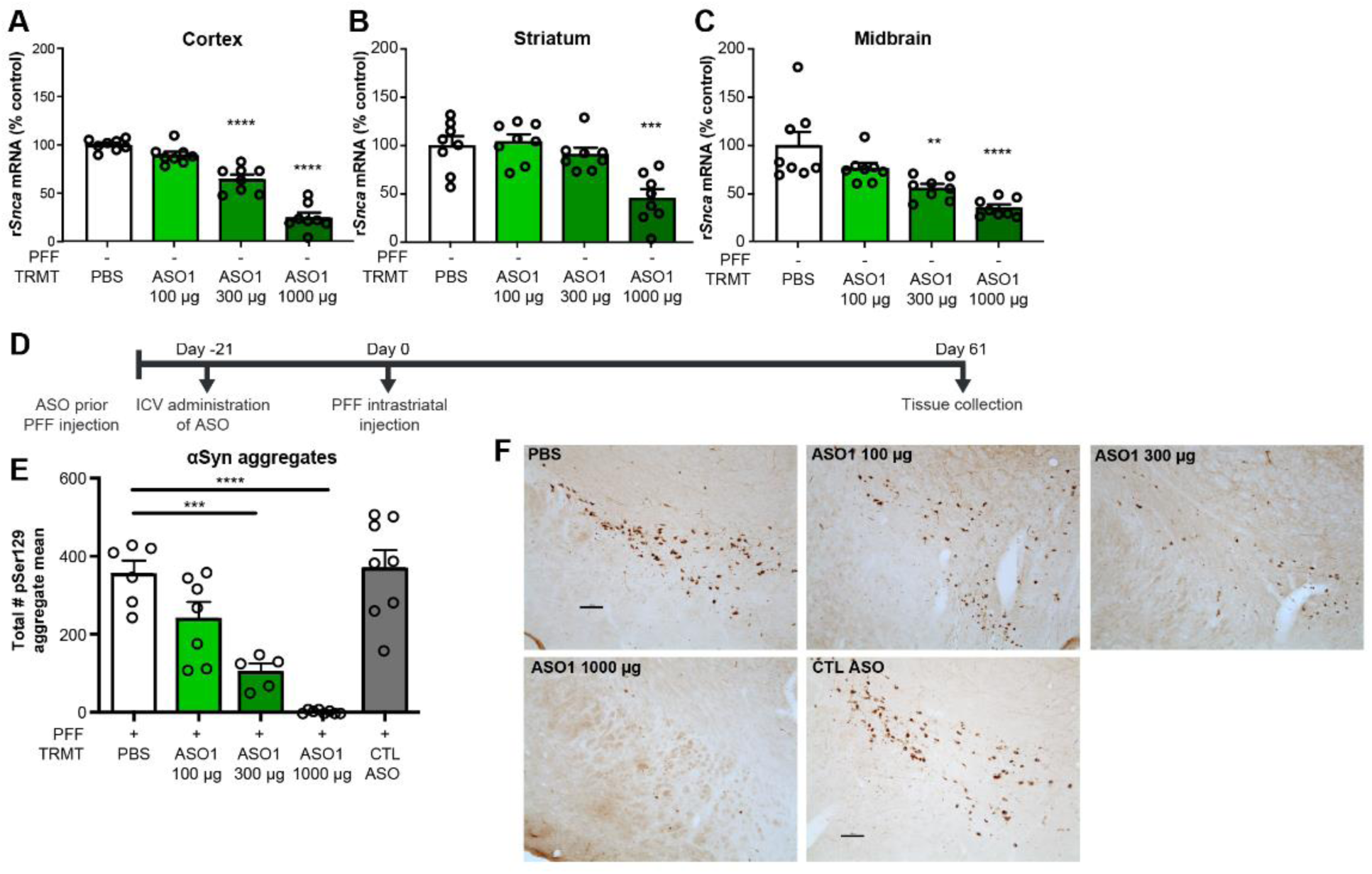
ASO-mediated reduction of *Snca* is dose-responsive, exhibits a prolonged duration of action, and dose-responsively prevents pathogenic aSyn aggregate deposition in an *in vivo* PFF model of PD. (**A** to **C**) 3 week dose response of rat *Snca* levels from cortical, striatal, and midbrain rat samples by RT-qPCR (n=8 per dose). (**D**) Timeline for ASO dose response administration prior to PFF injection paradigm. (**E**) Quantification of immunostaining for pSer129+ aggregate counts by total enumeration (n=6, 7, 5, 8, 8 for PBS, 100 µg, 300 µg, 1000 µg, and CTL ASO). (**F**) Representative images of pSer129+ aggregate counts in the substantia nigra. (Scale bar = 100 μm). Error bars represent ± s.e.m. *P<0.05, **P<0.01, ***P<0.001, ****P<0.0001 (Two-way ANOVA with Tukey post hoc analyses for duration of action with all other analyses using One-way ANOVA with Tukey post hoc analyses). PFF (pre-formed fibril), TRMT(treatment), CTL ASO (Control ASO)

A second *Snca* targeting ASO (ASO2) was also evaluated. *Snca* ASO2 also reduced pSer129+ aggregate counts, but to a lesser extent than ASO1 (fig. S1, F and G). This is consistent with ASO2 being a less potent molecule than ASO1 with almost no *Snca* mRNA suppression remaining 82 days post ASO administration (fig. S1, D and E), and more modest *Snca* mRNA suppression than ASO1 at 42 days post ASO administration (fig. S1, H to J). Taken together, these data suggest aSyn pathology is dependent on aSyn expression levels.

### ASO-mediated suppression of *Snca* prevents dopaminergic cell dysfunction in an *in vivo* PFF model of PD

PFF rodent models, like the human condition, undergo neurodegeneration in a number of CNS regions after pathology is established (Goedert et al., 2013; Luk et al., 2012; Paumier et al., 2015), this typically takes about 4-66 months after PFF injection. Fortunately, ASO1 exhibits a long duration of action with significant target suppression up to 84 days in cortex, striatum and midbrain then returning to baseline ∼160 days post a single ICV administration at 1000µg (Fig. 3A). To determine if ASO-mediated suppression of aSyn production could prevent TH loss, rats were administered a single ICV 1000µg dose of ASO1 prior to PFF injection and assessed 181 days later (Fig. 3B). ASO1 administration resulted in significant reduction of pSer129+ aggregates in comparison to PBS (∼53% reduction) and control ASO (∼48% reduction) treated rats (Fig. 3C), and significantly attenuated PFF-mediated TH loss in the SN compared to PBS at 181 days post a single ASO administration (Fig. 3D). Striatal dopamine levels were also normalized, in comparison to the respective contralateral side, with ASO1 treatment compared to PBS or control ASO (Fig. 3E). *Snca* mRNA had returned to normal levels at this 181 day timepoint (in comparison to PBS treated rats), which likely explains the modest accumulation of aSyn aggregates when compared to almost complete ablation of pathology at the 61 day timepoint when aSyn production was still suppressed (fig. S2, A and B in comparison to fig. S1, B, C, D, and E).

**Fig. 3.**
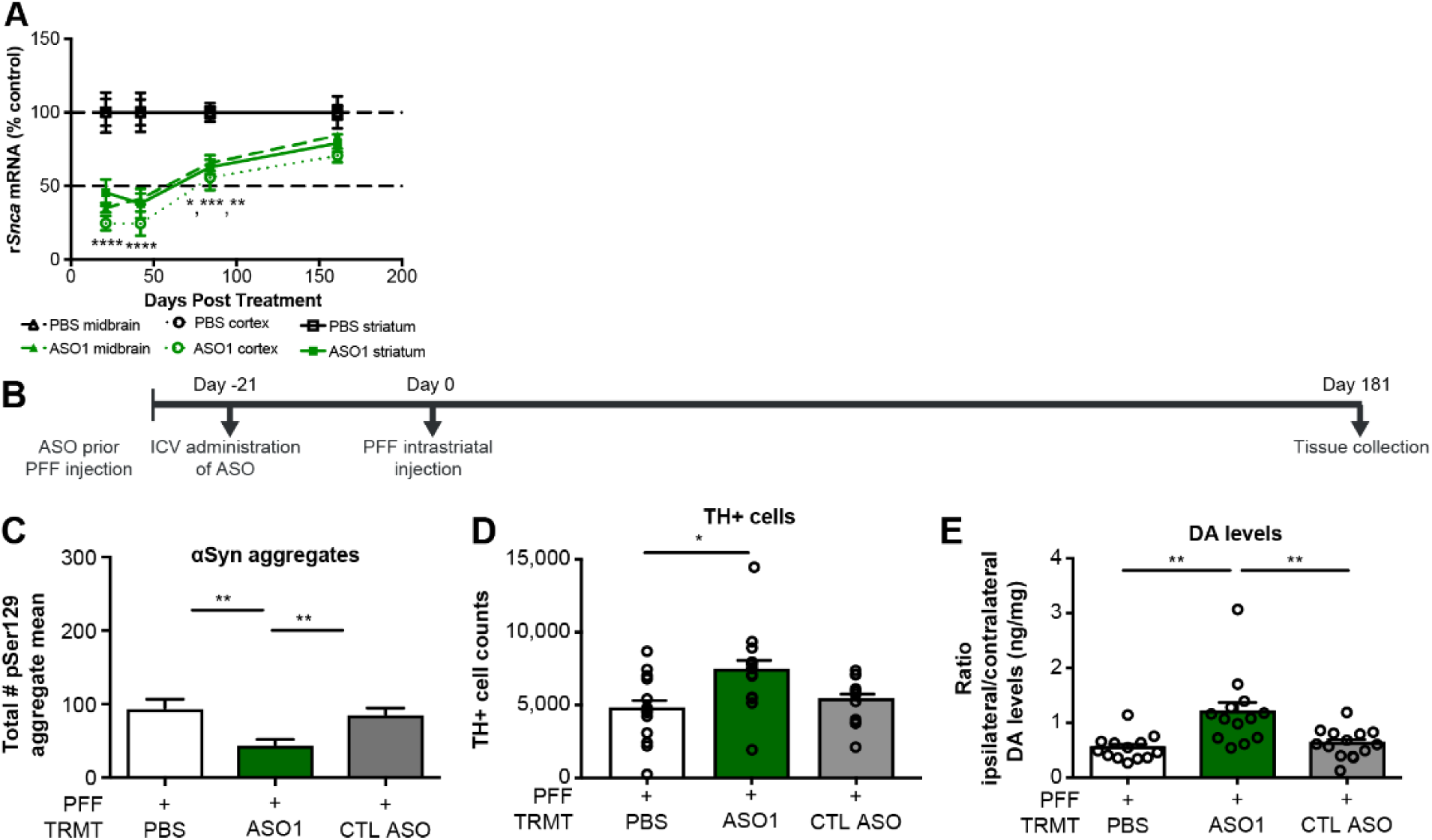
ASO-mediated suppression of *Snca* prevents dopaminergic cell dysfunction in an *in vivo* PFF model of PD. (**A**) Time course of *Snca* mRNA reduction (the same 1000 µg results for the three week time point in (**Fig. 2 A to C**) are included. (**B** to **E**) Results from ASO administration (1000 µg) prior to PFF injection paradigm in rats with study termination at 181 days. (**B**) Timeline for ASO administration prior to PFF injection paradigm in rats. (**C**) pSer129+ aggregate counts using total enumeration by IHC (n=13, 13, 15 for PBS, ASO1, and CTL ASO, respectively) (**D**) dopaminergic cell counts by IHC (by stereology) (n=13, 13, 12 for PBS, ASO1, and CTL ASO, respectively), and (**E**) striatal dopamine levels by HPLC (n=13, 13, 14 for PBS, ASO1, and CTL ASO, respectively) Data are ± s.e.m. *P<0.05, **P<0.001, ***P<0.0001, ****P<0.00001 (one-way ANOVA with Tukey post hoc analyses). PFF(pre-formed fibril), TRMT(treatment), CTL ASO(Control ASO).

### Pathogenic aSyn aggregate deposition is reversible and its amelioration reduces TH loss

To determine if ASO-mediated *Snca* suppression could be beneficial after established pathology, ASO1 was administered by ICV bolus at 700µg 14 days after PFF injection in mice, a timepoint with established pSer129+ aggregates in the SN (Fig. 4A). *Snca* mRNA was significantly reduced in the striatum and midbrain 56 days after PFF injection (Fig 4, B and C). Strikingly, ASO1 reduced the number of established aggregates (∼92% reduction, Fig 4D) 56 days post PFF injection. A trend toward improvement in a wire hang behavioral task was also found in mice (Fig. 4E). In addition, ASO1 was administered by ICV bolus at 1000µg 21 days after PFF injection in rats, a timepoint with established pSer129+ aggregates in the SN (Fig. 4, F and H). Though maximal PFF deposition occurs at 60 days post injection, rats euthanized 21 days post-PFF injection exhibit extensive aggregate deposition that was resolved by ASO administration ∼40 days later (Fig 4H). Strikingly, ASO1 reduced the number of established aggregates at 60 (∼90% reduction) and 81 (∼51% reduction) days post PFF injection (39 and 61 days post ASO administration) compared to pSer129+ aggregate counts at 21 days (PFF only). mRNA reduction was as expected (Fig. 4G). Similar reductions in pSer129+ aggregates were found in the insular cortex confirming widespread activity of the ASO (Fig. 4I). Thus, in both mice and rats ASO1 resulted in reversal of deposition of established aggregates.

**Fig. 4.**
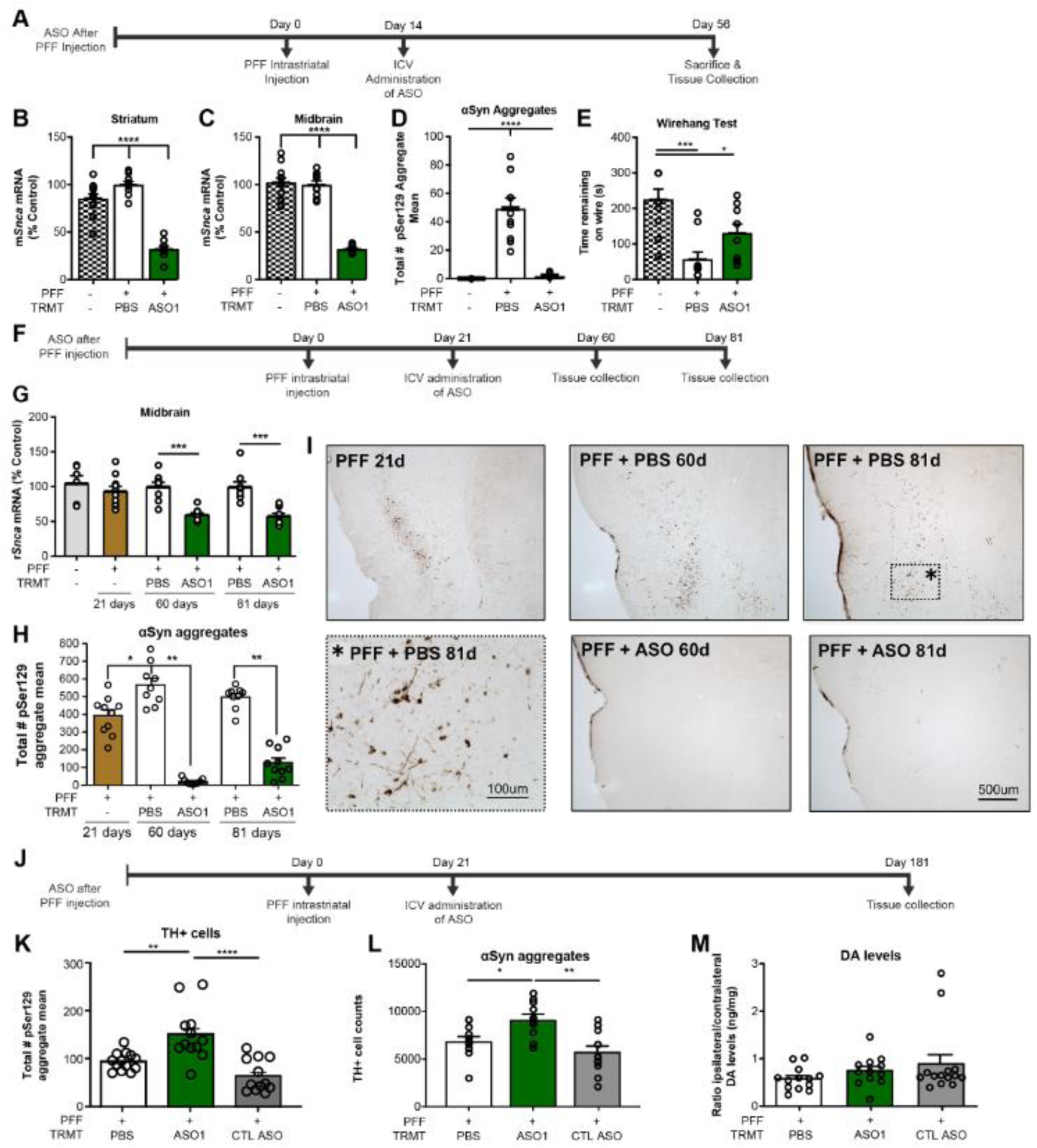
Pathogenic aSyn aggregate deposition is reversible and its amelioration reduces TH loss. (**A**) Timeline for ASO administration after PFF injection paradigm in the mouse. (**B** and **C**) mRNA reduction by RT-PCR, (n=12, 10, 10 for naïve, PBS, ASO1) (**D**) Quantification of aggregate reduction in the substantia nigra by IHC, (n=4, 10, 10 for naïve, PBS, and ASO1) and (**E**) performance on a wire hang task (n= 10, 10, 10 for naïve, PBS, ASO1, respectively). (**A** and **B**) Results from ASO administration after PFF injection with study termination at 60 and 81 days post PFF injection in the rat. (**F**) Timeline for ASO administration (1000 µg) for G to I. (**G**) Quantification of *Snca* mRNA reduction in the midbrain at each time point (**H**) Quantification of immunostaining for pSer129+ aggregate counts and (**I**) representative images from the insular cortex (n=10, 9, 9, 9, 10 for PFF only, PBS 60 days, ASO1 60 days, PBS 81 days, and ASO1 81 days (PBS at 60 days = 8). (**J** to **M**) Results from ASO administration (1000 µg) with study termination at 181 days. (**D**) Timeline for ASO administration for K to M. (**K**) TH+ cell counts by IHC (by stereology, (n=12, 11, 14 for PBS, ASO1, and CTL ASO, respectively) (**L**) Quantification of pSer129+ aggregate counts in the substantia nigra by IHC, (n=13, 12, 12 for PBS, ASO1, and CTL ASO, respectively) and (**M**) striatal dopamine levels by HPLC (n=13, 12, 14 for PBS, ASO1, and CTL ASO, respectively). Data are ± s.e.m. *P<0.05, **P<0.001, ***P<0.0001, ****P<0.00001. (one-way ANOVA with Tukey post hoc analyses). PFF(pre-formed fibril), TRMT(treatment), CTL ASO(Control ASO).

To determine if suppression of aSyn after established aggregate deposition could prevent TH loss, rats received a single 1000µg ICV injection of ASO1 administered 21 days after PFF injection and assessed 181 days post PFF (160 days post ASO administration) (Fig. 4J). ASO1 significantly attenuated PFF-mediated TH loss in the SN compared to PBS, while a control ASO did not (Fig. 4K). Interestingly, ASO1 administration resulted in a significant increase (∼22%) in pSer129+ aggregates compared to PBS and control ASO administered rats at 160 days post PFF injection (Fig. 4L), possibly due to reduced dysfunction of aggregate-bearing neurons. In this cohort, there were no significant differences in striatal dopamine levels for any of the treatment groups (Fig. 4M). The increase in aggregates in ASO1 administered animals relative to PBS 160 days after a single ASO administration is consistent with aSyn levels returning to normal over time (fig. S2, D and E), and likely higher than PBS due to the prevention of TH loss in ASO1 treated animals. Remarkably, the transient reduction in aSyn production after aggregates were established was sufficient to preserve TH expression in dopaminergic neurons highlighting the therapeutic value of directly targeting the underlying disease.

### Sustained reduction of *Snca* with ASO administered after aggregates are established reduces aggregate pathology and prevents TH loss

To evaluate a paradigm of sustained *Snca* reduction, mice were administered two ICV bolus 700µg administrations of ASO1 at 14 days prior to and 76 days after PFF injection, which resulted in sustained reduction in *Snca* mRNA with ∼50% reduction remaining 224 days after PFF injection and 238 days after the initial ASO administration (Fig. 5, A to C). Sustained *Snca* mRNA reduction resulted in sustained reduction in aSyn pathology and a prevention of TH loss 224 days after PFF injection (Fig. 5, D to F). *Snca* mRNA and pSer129+ aggregate reduction were similar to the ∼60 day mouse (Fig. 1 D to F) and rat studies (Fig. 2, A to G and fig. S1, B-K) though tissues were collected 224 days post PFF injection. Thus, maintaining *Snca* suppression prevented aggregates from accumulating and prevented TH loss.

**Fig. 5.**
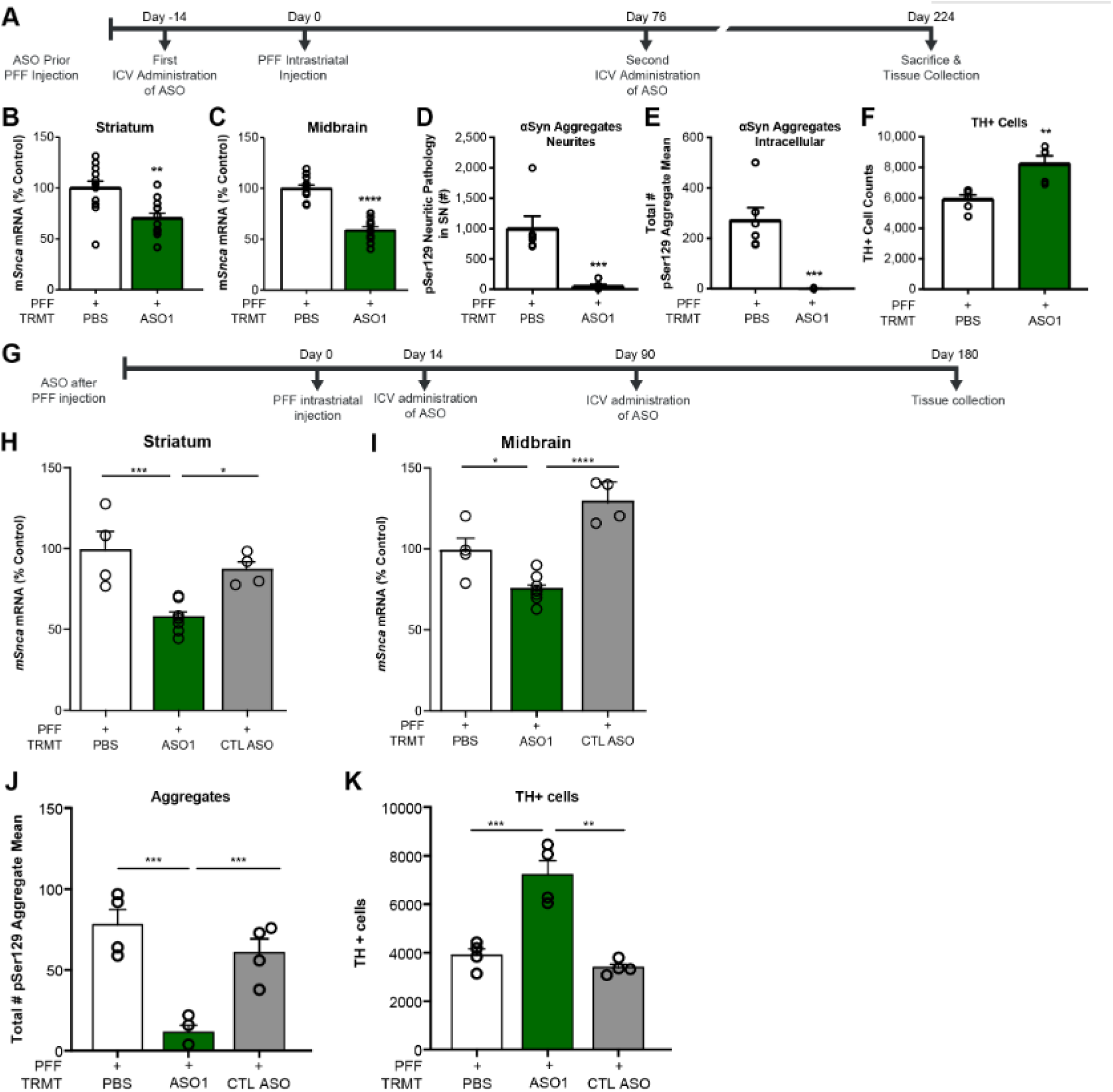
Sustained reduction of *Snca* with ASO administered after aggregates are established reduces aggregate pathology and prevents TH loss. (**A** to **F**) Results from ASO administration prior to PFF injection paradigm with sustained suppression and study termination at 224 days post PFF injection in mice. (**A**) Timeline for ASO administration (700µg) prior to PFF injection paradigm. (**B** and **C**) mRNA reduction in the striatum and midbrain. pSer129+ aggregate counts quantified by IHC in (**D**) neurites and (**E**) cell bodies. (**F**) Dopaminergic cell counts quantified by IHC. (n=12 for PBS and ASO1) (n=6 for immunohistochemistry endpoints). (**G** to **K**) Results from ASO administration after PFF injection paradigm with sustained suppression and study termination at 224 days post PFF injection in mice. (700 µg). (**G**) Timeline for ASO administration (700µg) after to PFF injection paradigm. (**H** and **I**) mRNA reduction in the striatum and midbrain. pSer129+ aggregate counts quantified by IHC in (**J**). (**K**) Dopaminergic cell counts quantified by IHC. (n=4). Data are ± s.e.m. *P<0.05, **P<0.001, ***P<0.0001, ****P<0.00001. (one-way ANOVA with Tukey post hoc analyses). PFF(pre-formed fibril), TRMT(treatment), CTL ASO(Control ASO).

To determine if prolonged ASO-mediated *Snca* suppression could be more beneficial than transient reduction after established pathology, ASO1 was administered ICV at 14 and 90 days following PFF injection with study termination at 180 days post PFF in mice (Fig. 5G). *Snca* mRNA and PSer129+ aggregates were reduced in the substantia nigra (Fig. 5, H to J). TH positive loss was also prevented with prolonged aSyn reduction post PFF administration in comparison to PBS administered mice (Fig. 5K). This differs from single ICV dose injection following PFF in which aggregate number was increased in ASO1 treated rats in comparison to PBS treated rats 180 days following PFF injection when mRNA expression had returned to baseline levels (Fig. 4L and Fig. 3A) Thus, therapeutic treatment with *Snca* ASOs administered with established aggregates requires sustained reduction of *Snca* to prevent aggregates from accumulating once *Snca* RNA levels return to baseline.

### Human SNCA ASOs are potent, suppress aSyn broadly in the primate CNS and CSF aSyn is a potential pharmacodynamic biomarker

The ASOs used thus far target the rodent *Snca* transcripts. To treat human patients, we designed ASOs to suppress the human *SNCA* transcript, using similar designs and chemistries as ASOs already in clinical testing (Tabrizi et al., 2019). Human sequence targeting *SNCA* ASOs hASO1 and hASO2 (Table 1B) suppress human *SNCA* mRNA in a dose- and concentration-dependent manner (Fig. 6, A to D and fig. S3, A and B) *in vitro* in SH-SY5Y human cells and *in vivo* in the aSyn wild type human full-length transgenic mouse (similar to (Nussbaum and Ellis, 2003). The human *SNCA* ASOs also exhibit an extended duration of action lasting 10 weeks after a single administration (fig. S3, C to E). To determine the activity and distribution of human ASOs in a larger brain, hASO1 and hASO2 were dose intrathecally (IT) in non-human primates (NHPs, cynomolgus). Following repeated IT delivery of hASO1 or hASO2 to the NHPs, *SNCA* mRNA (RT-qPCR) and aSyn protein (ELISA) are reduced throughout the brain and spinal cord (Fig. 6, E to H). Further analyses of NHP brain tissue by *in situ* hybridization (*SNCA* mRNA) and immunohistochemistry (ASO and aSyn protein) support the conclusion that *SNCA* ASOs distribute broadly and result in reduction of *SNCA* mRNA and protein throughout the brain and spinal cord (Fig. 6I and fig. S4), including regions implicated in PD. aSyn protein in the CSF is significantly reduced with hASO1 in comparison to vehicle (aCSF) administered NHPs (Fig. 6J). In addition, aSyn protein reduction in the frontal cortex correlates to aSyn protein reduction in the CSF, suggesting the use of aSyn in the CSF as a potential pharmacodynamic biomarker (Fig. 6K).

**Fig. 6.**
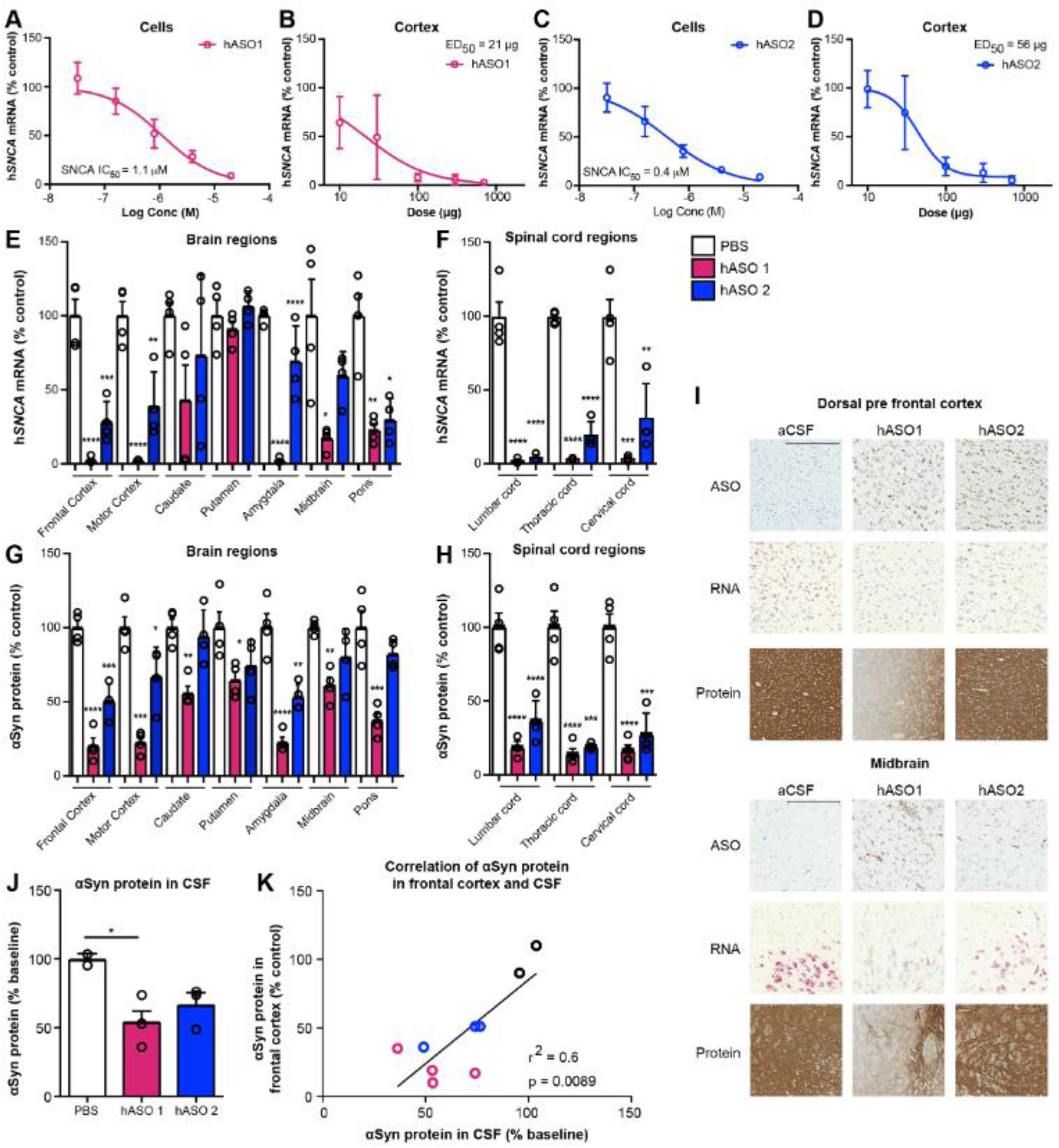
Human *SNCA* ASOs are potent, suppress aSyn broadly in the primate CNS and CSF aSyn is a suitable pharmacodynamic biomarker. *In vitro* dose response of human *SNCA* hASO1 and hASO2 in (**A** and **C**) SHSY5Y cells and (**B** and **D**) *in vivo* dose response in SNCA-PAC mouse cortex 2 weeks post injection (n=10 except 700 µg n=7 for hASO1) (n=8 for for hASO2 (2 mice removed with ROUT analysis from 300 µg and 700 µg groups, combination of two studies of n=4 each, n=12 for all PBS groups). (**E** and **F**) mRNA by RT-PCR and (**G** and **H**) protein by ELISA throughout the brain and spinal cord of the NHP (n=4 for PBS, hASO1, and hASO2). (**I**) Representative images for IHC (ASO and aSyn antibody) or ISH (*SNCA* mRNA). Scale bar = 600 µm for protein/300 µm for ASO and ISH results. (**J**) Quantification of aSyn protein by ELISA in CSF with (**K**) correlation of aSyn protein levels in the frontal cortex and CSF (n=2, 3, and 4 for PBS, hASO1, and hASO2). Data are ± s.e.m except for SHSY5Y, which are StDev. *P<0.05, **P<0.001, ***P<0.0001, ****P<0.00001. (One-way ANOVA with Tukey post hoc analyses).

## Discussion

Our study demonstrates that ASO-mediated suppression of *Snca* prevented and reversed the progression of aSyn mediated pathology in rodent transmission models of PD demonstrating the potential of *SNCA* ASOs as a therapy for PD patients. Long term studies with sustained reduction of aSyn prevented and even delayed pathology and associated TH loss. Furthermore, central delivery of human *SNCA* ASOs reduced expression of mRNA and protein throughout the brains of both the humanized mouse and NHPs demonstrating that human *SNCA* ASOs are active in regions of the brain susceptible to PD in a larger species. In NHPs the reduction of aSyn was reflected in the CSF supporting further investigation of the use of aSyn protein levels in the CSF as a target engagement biomarker to allow evaluation of efficacy of human *SNCA* ASOs in the clinic.

PD and other synucleinopathies are generally characterized as toxic gain of function in *SNCA*, suggesting that therapeutics designed to lower aSyn production would be beneficial to patients (Giasson et al., 2002; Klein et al., 2002; Kruger et al., 1998; Zarranz et al., 2004). To date, there are multiple protein-based therapeutic strategies being investigated targeting different forms and conformational states of aSyn in clinical trials (Kingwell, 2017). However, the toxic species of aSyn has not yet been confirmed, potentially limiting therapeutic benefit of these approaches (Dehay et al., 2015; Kingwell, 2017). In addition, protein targeting therapies hypothesized to clear aSyn aggregates as they are transmitted from cell to cell may not be as effective in targeting intracellular Lewy body inclusions. An ASO therapy can potentially overcome these limitations by targeting mRNA intracellularly and reducing all forms of aSyn protein (Bennett and Swayze, 2010; Rigo et al., 2014). Indeed, ASO-mediated suppression of aSyn reduced aSyn pathology in a dose dependent manner. In all cases, the levels of aSyn pathology corresponded to the levels of endogenous aSyn production. An ASO less effective at suppressing *Snca* mRNA was less effective at suppressing aSyn pathology. If the *Snca* mRNA levels were allowed to return to normal levels, the aSyn pathology returned, however, this increase was prevented with continued suppression of aSyn production. Taken together, these data illustrate the link between production of endogenous aSyn and aSyn pathology.

Our observation that ASO treatment was beneficial in reducing the number of pSyn aggregates at a time when pre-existing pathology is present is particularly interesting. This is consistent with previous work reporting reversal of deposition of soluble aSyn as well as insoluble forms of pathology following reduction of aSyn production by tetracycline-controlled transactivator (tTA) to prevent expression of the A53T mutant aSyn transgene (Lim et al., 2011). In the genetic study, stopping production also reversed detrimental changes in hippocampal synaptic markers, and hippocampal memory deficits (Lim et al., 2011). Our results replicate and extend these findings with a therapeutically relevant modality, and in a model where we are targeting endogenous aSyn, rather than a transgene. In both cases, stopping aSyn production had a dramatic effect on aSyn pathology and neuronal health. This finding may suggest that *de novo* production and recruitment of aSyn is required for maintaining the stability of aggregates until a threshold is reached for irreversible cell death. This concept remains to be explored. Regardless, these data suggest that treatment initiation after disease onset in sporadic patients has the potential to reverse disease.

Many toxin, genetic, viral-mediated, and alpha-synuclein injection Parkinson disease animal models have been generated to evaluate the role of aSyn in Parkinson’s disease (Jagmag et al., 2015; Koprich et al., 2017; Recasens et al., 2014). Each model exhibits advantages and limitations (Koprich et al., 2017). The aSyn PFF injection models, used here, exhibits advantages in modeling sporadic PD by relying on fibril seeding and templating of normal endogenous levels of aSyn in specific interconnected brain circuits, avoiding the global overexpression produced in germline transgenic models and the forced local supraphysiological overexpression common in viral vector models (Duffy et al., 2018). The evolution of aggregate formation in morphology (diffuse punctate to compact) and important characteristics of Lewy bodies (e.g., proteinase-K resistance, Thioflavin-S positive) progressing over time to result in DA neuron degeneration at 6 months post-PFF injection, provides a platform for analyzing the potential of neuroprotective therapies, such as ASOs, for translation to treatment of idiopathic PD. In this regard, our findings are promising.

Few tolerability concerns exist for lowering *SNCA* as evidenced by genetically engineered *Snca* deficient mice (Abeliovich et al., 2000; Cabin et al., 2002; Chandra et al., 2004; Goldberg and Lansbury, 2000; Greten-Harrison et al., 2010). There are reports of cell death *in vivo* using an shRNA against Snca (Benskey et al., 2018; Collier et al., 2016), which was not found here with ASOs or in other publications using siRNAs, shRNAs, or ASOs against *Snca* (Alarcon-Aris et al., 2018; Uehara et al., 2019; Zharikov et al., 2015). The aSyn antibodies, Prasinezumab (PRX002) and Cinpanemab (BIIB054), completed first-in-human trials in which no serious adverse events were found with aSyn lowering (Brys et al., 2019; Schenk et al., 2017). In addition, our animal model studies and others (Games et al., 2014; Spencer et al., 2017; Tran et al., 2014; Zharikov et al., 2015) have shown a benefit with less than 50% reduction of *Snca* mRNA indicating that only a partial reduction of *SNCA* will be needed for therapeutic benefit, as mentioned above.

aSyn protein exhibits a profile desirable for a pharmacodynamic biomarker as reduction in aSyn protein in the CSF correlated with aSyn protein reduction in the frontal cortex in the NHP. aSyn protein is measurable in human CSF, has been shown in patients to not correlate with clinical progression, and levels are relatively stable over 24 months, thus suggesting aSyn levels in the CSF may potentially be used in clinical trials as a pharmacodynamic marker to support demonstration of efficacy of aSyn modulating therapies (Dolatshahi et al., 2018; Mollenhauer, 2014; Mollenhauer et al., 2017; Parnetti et al., 2016; Zhou et al., 2015). Continued evaluation and improved methods for detecting aSyn in the CSF and other fluids are warranted, such as electrochemiluminescense-based detection (Kruse and Mollenhauer, 2019).

ASOs represent a therapeutic approach for directly lowering *SNCA* production because ASOs are sequence-specific and can reach central nervous system targets by intrathecal delivery. The ASO platform is quickly becoming a realistic therapeutic strategy for the treatment of central nervous system (CNS) diseases due partly to recent advances in ASO design which have improved stability, affinity, and potency, as well as improved tolerability (Bennett and Swayze, 2010; Rigo et al., 2014). ASOs have also been shown to distribute widely within the brain and spinal cord in the NHP shown here and elsewhere (Kordasiewicz et al., 2012; Lagier-Tourenne et al., 2013; Passini et al., 2011) in addition to exhibiting a long duration of effect shown here and elsewhere (Friedrich et al., 2018; Hagemann et al., 2018; McLoughlin et al., 2018; Scoles et al., 2017; Zhao et al., 2017). Feasibility of the approach is supported by FDA approval of Spinraza© for the treatment of spinal muscular atrophy (Chiriboga et al., 2016; Finkel et al., 2016), the recently completed clinical trial with an ASO for Huntingtin’s (Htt) disease (Rodrigues and Wild, 2018; Tabrizi et al., 2019), and ongoing trials for a SOD1-targeted ASO for amyotrophic lateral sclerosis (NCT02623699), a C9ORF72 ASO for ALS (NCT03626012), a LRRK2-targeted ASO for Parkinson’s disease (NCT03976349), and a MAPT targeting ASO therapy for Alzheimer’s disease (NCT03186989). To explore the translational potential of an *SNCA* ASO therapy, we examined target engagement of human *SNCA* targeting ASOs in a human cell line, a human aSyn transgenic mouse model and NHPs. The *SNCA* targeting molecules described here have sufficient potency and duration of action for use in human patients. Thus, ASOs designed against human *SNCA* have the potential to be a disease-modifying therapeutic for PD patients.

## Materials and Methods

### Oligonucleotide synthesis

The synthesis and purification of all lyophilized ASOs was formulated in PBS without Ca/Mg (Gibco: 14190) as previously described and stored at −20°C (Ostergaard et al., 2013; Seth et al., 2010). Sequences and chemistries used are listed (fig. S1A and B).

### Cell culture PFF experiments

Timed-pregnant CD1 mice (Charles River Laboratories) were utilized for primary neuronal cultures. Hippocampal neurons were prepared from embryos (E16-18) as previously described (Volpicelli-Daley et al., 2014). All other methods including PFF generation, PFF treatment, and ASO addition to cultures were performed as previously described (Luna et al., 2018).

PSer129+ pathology and double strand DNA breaks by γH2AX Ser139 were quantified as previously described (Tapias et al., 2017).

### Rats

Adult male Sprague Dawley rats (200-225g; Harlan Laboratory, Indianapolis, IN) were utilized in rat experiments. Rat PFF studies were conducted at Michigan State University (MSU) and rat and mouse studies were conducted at Ionis pharmaceuticals. Rats at MSU were housed in the Van Andel Research Institute vivarium and mice and rats at Ionis pharmaceuticals were housed in the vivarium. Both animal facilities are accredited by the Association for the Assessment and Accreditation of Laboratory Animal Care and complied with all Federal animal care and use guidelines for both institutions. The Institutional Animal Care and Use Committee approved all protocols for both institutions.

### Intrastriatal PFF injections in rats

Purification of recombinant *in vitro* fibril assembly was performed as previously described (Paumier et al., 2015). Animals were monitored weekly following surgery and sacrificed at various time points.

### ICV injection in rats

Rats were injected into the right cerebroventricle (ICV) using a stereotactic device. For ICV bolus injections the coordinates were −1.0 mm anterior/posterior and 1.5 mm to the right medial/lateral are used. The needle is lowered −3.7mm dorsal/ventral. The proper amount of injection solution (30 μL) was injected by hand at injection rates of approximately 1 μL/second with a 5-minute wait following completion of injection. The incision was sutured closed using one horizontal mattress stitch with 3-*O* Ethilon suture. The animals were then allowed to recover from the anesthesia in their home cage.

### Tissue processing in rats

Following in-life completion rats were euthanized by CO_2_ asphyxiation. 2 mm sections of spinal cord and different brain regions were collected for mRNA analysis. Brains perfused with physiological saline were cooled in iced saline and cut coronally in a 2 mm thick slab immediately rostral to the hypothalamus. The section was placed onto a petri dish on ice and further dissected into 3 pieces each for striatum and overlying cortex. For striatum: dorsal lateral striatum for HPLC, dorsal intermediate striatum for mRNA, and dorsal medial striatum for protein. For cortex: medial for HPLC, intermediate for mRNA, lateral for protein. Dissections were optimized for HPLC analysis corresponding to the pattern of dopamine innervation. Each dissected region was frozen at −80 until analysis. Immunohistochemistry/ immunofluorescence/ pSer129+ aggregate counts (total enumeration)/stereology were performed as previously described (Paumier et al., 2015).

### Immunohistochemistry in rats

Used primary mouse anti-pSyn (81a-1:10,000 (Luk et al., 2012)), mouse-anti-TH (1:8000; Immunostar, Hudson, WI), antibodies overnight at 4°C. Then, sections were incubated in biotinylated secondary antisera against either mouse (1:400, Millipore, Temecula, CA) or rabbit IgG (1:400, Millipore, Temecula, CA) followed by Vector ABC detection kit (Vector Labs, Burlingame, CA). Antibody labeling was visualized by exposure to 0.5 mg/ml 3,3’ diaminobenzidine (DAB) and 0.03% H_2_O_2_ in Tris buffer. Sections were mounted on subbed slides, dehydrated to xylene and coverslipped with Cytoseal (Richard-Allan Scientific, Waltham, MA).

### Immunofluorescence in rats

For DAPI staining, an additional three-minute incubation in Tris buffer with DAPI (1:500; Invitrogen, Carlsbad, CA) was performed. Primary antibodies used include rabbit anti-TH (1:4000; Millipore, Temecula, CA), mouse anti-pSyn (81a-1:15,000),. Secondary antibodies used include Alexa Fluor 568 goat anti-mouse IgG (1:500; Invitrogen, Carlsbad, CA) and Alexa Fluor 488 goat anti-rabbit IgG (1:500; Invitrogen, Carlsbad, CA).

### Medial Terminal Nucleus (MTN) counts of TH neurons in the SN in rats

TH neurons from three sections of the SN, easily identified by proximity to the medial terminal nucleus of the accessory optic tract (−5.04 mm, −5.28 mm and −5.52 mm relative to bregma), were quantified as previously described (Gombash et al., 2014). MicroBrightfield stereological software (MBF Bioscience, Williston, VT) was used to assess the total number of aggregates (defined as dense, darkly stained cores of phosphorylated aSyn staining) within the SN at various time points. Using a Nikon Eclipse 80i microscope, Retiga 4000R (QImaging, Surrey, BC, Canada) and the Microbrightfield StereoInvestigator software (Microbrightfield Bioscience, Burlingame, Virginia, USA), MTN neuron quantification was completed by drawing a contour around the SN borders using a 4X objective. Virtual markers were then placed on TH+ neurons at a 20X objective and quantified. Total TH neuron numbers in both the ipsilateral and contralateral SN were averaged for the three medial terminal nucleus sections counted. MTN counts were used for fig. S2, C. Differences in n’s within an experiment are due to technical reasons.

### Stereology in rats

MicroBrightfield stereological software (MBF Bioscience, Williston, VT) was used to assess total population cell counts in the substantia nigra pars compacta (SNpc). The total number of stained neurons was calculated using optical fractionator estimations and the variability within animals was assessed via the Gundersen Coefficient of Error (< 0.1) (Gundersen et al., 1999).

### High performance liquid chromatography (HPLC) in rats

A dorsolateral striatal tissue punch was taken from both hemispheres. Frozen punches were placed individually in vials supercooled on dry ice and stored at −80 °C until analysis. Tissue was homogenized and analyzed as described previously (Koprich et al., 2003a; Koprich et al., 2003b). The Pierce BCA Protein Kit (Rockford, IL) was utilized for protein determination. Samples were separated on a Microsorb MV C-18 column (5 Am, 4.6–250 mm, Varian, Palo Alto, CA) and simultaneously examined for norepinephrine, serotonin, dopamine, 3,4-dihydroxyphenylacetic acid (DOPAC) and homovanillic acid (HVA). Compounds were detected using a 12-channel coulometric array detector (CoulArray 5200, ESA, Chelmsford, MA) attached to a Waters 2695 Solvent Delivery System (Waters, Milford, MA) under the following conditions: flow rate of 1 ml/min; detection potentials of 50, 175, 350, 400 and 525 mV; and scrubbing potential of 650 mV. The mobile phase consisted of a 10% methanol solution in distilled H2O containing 21 g/l (0.1 M) citric acid, 10.65 g/l (0.075 M) Na2HPO4, 176 mg/l (0.8 M) heptanesulfonic acid and 36 mg/l (0.097 mM) EDTA at a pH of 4.1.

### Mice

Adult male and female mice (20-32 g;Taconic Biosciences, Hudson, NY) were utilized in all experiments. All mouse studies using either C57/Bl6 or human wild type SNCA-PAC (Licensed from Mayo Foundation for Medical Education and Research) mice were conducted at Ionis pharmaceuticals. The Institutional Animal Care and Use Committee approved all protocols.

### Intrastriatal PFF injections in mice

Mouse PFF studies were performed as described previously including pSer129+ aggregate counts, dopaminergic cell counts, and wire hang task in mice (Zhao et al., 2017). Purification of recombinant *in vitro* fibril assembly was performed as previously described (Luk et al., 2012; Volpicelli-Daley et al., 2014; Zhao et al., 2017). Mice were injected in the striatum with 5 µg of aSyn PFFs in 2 µL of dosing solution.

### ICV ASO injection in mice

Mice were injected into the right cerebroventricle (ICV) using a stereotactic device as previously described (Zhao et al., 2017). For ICV bolus injections the coordinates 0.3 mm anterior to bregma, 1.0 mm right lateral, and −3.0 mm ventral were used. 10μL of injection solution was injected by hand at injection rates of approximately 1 μL/second with a 5-minute wait following completion of injection. The incision was sutured closed using one horizontal mattress stitch with 5-*O* Ethilon suture. The animals were then allowed to recover from the anesthesia in their home cage.

### Non-human primates

The non-human primate (cynomolgus) study was performed at Covance laboratories GmbH. Covance Laboratories GmbH test facility is fully accredited by the AAALAC. All procedures in this study plan are in compliance with the German Animal Welfare Act and are approved by the local IACUC, and performed as described previously (DeVos et al., 2017). NHPs (n=4) were administered, by intrathecal bolus injection, 5 doses of ASO or aCSF on Day 1, 14, 28, 56, and 84 with 1.0 mL dosing volume using a 35 mg/mL dosing solution and euthanized on day 91.

### aSyn enzyme-linked immunosorbent assay (ELISA)

Roughly 50 mg of NHP brain tissue was added to 1 ml of RIPA buffer (Boston Bioproducts) with protease and phosphatase inhibitor tablets (Sigma-Aldritch) in a 2 ml Lysing Matrix D tube (MP Biomedicals). The samples were homogenized using an MP Fastprep-24 (MP Biomedicals), and total protein was quantified using the Pierce BCA Protein Assay kit (Thermo Fisher) and normalized to 1 mg/ml. Tissue samples were diluted either 1:100 (spinal cord), 1:2000 (brain tissue) or 1:10 (CSF) prior to alpha synuclein protein determination. Alpha synuclein protein concentrations were measured using the LEGENDMAX Human Alpha Synuclein ELISA Kit (Biolegend) following the manufacturer’s protocol. CSF hemoglobin levels were also analyzed using the human hemoglobin ELISA kit (Bethyl Laboratories). This was done to assess the impact of blood contamination on the alpha synuclein levels detected in CSF due to the high levels of alpha synuclein contained in red blood cells. Samples with greater than 1000 ng/ml HgB were excluded from the CSF analysis.

### Antisense oligonucleotide immunohistochemistry in NHP tissue

Immunohistochemical staining was performed on a Ventana *DISCOVERY ULTRA* autostainer (Ventana). Staining was done according to the manufacturer’s instructions. Briefly, five micrometer thick sections were deparaffinized and rehydrated then subjected to antigen retrieval using Proteinase K (Dako) for 4 minutes, followed by incubation with a Rabbit polyclonal anti-ASO antibody at 1: 10,000 dilution (Ionis, #ASO 6651) for 60 minutes. The signal was detected with polymer-based secondary antibody-Discovery OmniMap anti-Rb HRP and Discovery ChromoMap DAB kits (Ventana). Control samples were also stained with rabbit IgG (Cell signaling Technologies, #2792) as isotype controls. All slides were scanned on a 3DHISTECH Panoramic P250 Slide Scanner. ASO images were scanned at 20X magnification. All images were taken as screen shots from virtual slides. All images were analyzed with custom image analysis algorithms performed on the VisioPharm software platform. Target brain regions on each slide were manually outlined in Visiopharm to designate areas for algorithm analysis. In the VisioPharm analysis algorithm, ASO staining was assessed as positively stained region area broken down into three levels of intensity. A minimum threshold for ASO staining was determined by assessing background levels of DAB (brown) staining. A maximum threshold of 100% intensity of DAB staining was used. ASO staining ranges were determined independently per stain. This range was evenly divided into low, medium, and high intensity bins, each area of staining was calculated as a percentage of the total outlined tissue area (brain region).

### SNCA-RNA *in situ* hybridization (ISH) staining in NHP tissue

ISH staining was performed on a Leica Biosystems’ BOND RX Autostainer (Leica Biosystems) using mRNAscope^®^ 2.5 LS Reagent Kit—RED (Advanced Cell Diagnostics). Staining was done according to the manufacturer’s instructions. Briefly, five micrometer thick sections were deparaffinized and rehydrated then subjected to pretreatment, followed by specific probe-SNCA (ACD, #421318) hybridized to target mRNA. The signal was amplified using multiple steps, followed by hybridization to alkaline phosphatase (AP)-labeled probes and detection using Fast Red (ACD, #322150). The tissue quality was examined by positive control probe-PPIB (ACD, #424148), and background staining was assessed by negative control probe-dapB (ACD, #312038). All slides were scanned on a 3DHISTECH Panoramic P250 Slide Scanner. ISH images were scanned at 40X magnification. All images were taken as screen shots from virtual slides. All images were analyzed with custom image analysis algorithms performed on the VisioPharm software platform. Target brain regions on each slide were manually outlined in Visiopharm to designate areas for algorithm analysis. In the VisioPharm analysis algorithm, aSyn mRNA counts were determined by detecting and counting SNCA ISH labeled dots within each region. These counts were normalized by the total outlined tissue area (brain region) in mm^2^.

### aSyn immunohistochemistry in NHP tissue

Immunohistochemical staining was performed on a Ventana *DISCOVERY ULTRA* autostainer (Ventana). Staining was done according to the manufacturer’s instructions. Briefly, five micrometer thick sections were deparaffinized and rehydrated then subjected to antigen retrieval using Ventana CC1 (EDTA pH 8) for 64 minutes, followed by incubation with a Mouse monoclonal antibody aSyn211 (Santa Cruz, #SC-12767) at 0.25 μg/ml for 60 minutes. The signal was detected with polymer-based secondary antibody-Discovery OmniMap anti-Ms HRP for 16 minutes. Control samples were also stained with mouse IgG (Cell signaling Technologies) as isotype controls. All slides were scanned on a 3DHISTECH Panoramic P250 Slide Scanner. aSyn protein images were scanned at 20X magnification. All images were taken as screen shots from virtual slides. All images were analyzed with custom image analysis algorithms performed on the VisioPharm software platform. Target brain regions on each slide were manually outlined in Visiopharm to designate areas for algorithm analysis. In the VisioPharm analysis algorithm, aSyn Protein staining was assessed as positively stained region area broken down into three levels of intensity. A minimum threshold for aSyn protein was determined by assessing background levels of DAB (brown) staining. A maximum threshold of 100% intensity of DAB staining was used. aSyn Protein ranges were determined independently per stain. This range was evenly divided into low, medium, and high intensity bins, each area of staining was calculated as a percentage of the total outlined tissue area (brain region).

### mRNA purification and analysis for mice, rats, and non-human primates

Approximately 10 mg of tissue per sample was homogenized in guanidinium isothiocynate. Total mRNA was purified further using a mini-mRNA purification kit (Qiagen, Valencia, CA). After quantitation, the tissues were subjected to real time PCR analysis. The Life Technologies ABI StepOne Plus Sequence Detection System was employed. Briefly, 20 µl RT-PCR reactions containing 5µl of mRNA were run with the RNeasy 96 kit reagents and the primer probe sets listed in the materials section optimized from manufacturer’s instructions. The following sequences of primers and probes were used: mouse *Snca* 5’-GTCATTGCACCCAATCTCCTAAG-3’ (forward), 5’-GACTGGGCACATTGGAACTGA-3’ (reverse), and 5’-FAM-CGGCTGCTCTTCCATGGCGTACAAX-TAMRA-3’ (probe); rat *Snca* 5’-GATGGGCAAGGGTGAAGAAG-3’ (forward), 5’-GCTAGGGTCCACAGGCATGT-3’ (reverse), and 5’-FAM-TACCCACAAGAGGGAAT-MGB-3’ (probe) human *SNCA* 5’-TGGCAGAAGCAGCAGGAAA-3’ (forward), 5’-TCCTTGGTTTTGGAGCCTACA-3’ (reverse), and 5’-FAM-CAAAAGAGGGTGTTCTC-TAMRA-3’ (probe); mouse *cyclophilin A* 5’-TCGCCGCTTGCTGCA-3’ (forward), 5’-ATCGGCCGTGATGTCGA-3’ (reverse), and 5’-FAM-CCATGGTCAACCCCACCGTGTTCX-TAMRA-3’; Target *SNCA* mRNA was then normalized to Cyclophilin A mRNA levels from the same mRNA sample. SNCA mRNA from ASO treated animals was further normalized to the group mean of PBS treated animals, and expressed as percent control. All qPCR reactions were run in triplicate. Data were analyzed using Microsoft Excel (v14.4).

### SH-SY5Y Cell Culture Assay

SH-SY5Y cells (CRL-2266, ATCC) were cultured in growth medium at 37° C and 10% CO2. ASO was electroporated into cells, by pulsing once at 160V for 6mS with the ECM 830 instrument (Harvard Apparatus). After 24 hr, the cells were washed 1X with PBS before lysing for mRNA isolation and analysis. For each treatment condition quadruplicate wells were tested.

### RNA purification and analysis for cell culture

The mRNA was purified with a glass fiber filter plate (Pall # 5072) and chaotropic salts. The human *SNCA* message level was quantitated with RT-qPCR on the QS7 instrument (Applied Biosystems). Total mRNA levels were measured with the Quant-iT™ RiboGreen^®^ mRNA reagent and used to normalize the *SNCA* data. Data were analyzed using Microsoft Excel (v14.4) and GraphPad Prism (v6).

### Statistical Analysis

Statistical tests were completed using either GraphPad Prism software (version 6, GraphPad, La Jolla, CA) or Microsoft excel (v14.4). Data are expressed as mean +/− s.e.m., unless otherwise noted. For two group comparisons t-tests were used. For more than two group comparisons, one-way ANOVAs were used. Comparisons of multiple groups made across time points were analyzed using a two-way ANOVA. When appropriate, post-hoc comparisons were made between groups using Tukey or Bonferonni post hoc tests. A ROUT analysis was performed to determine outliers using (Q=1%). The level of significance was set at P ≤ 0.05. Tests used are reported in the figure legends.

## Summary of supplemental material

Supplemental material consists of ASO sequences, confirmation of POC results with an additional *Snca* ASO, non-critical RNA results, additional characterization of human *SNCA* ASOs in transgenic mice, and supporting histological evidence from NHP study.

## Author contributions

The study was conceived by T.A.C. and H.B.K. Experiments were designed by T.A.C., H.B.K., H.Z, and K.L.P. ASOs were designed by T.A.C, H.B.K, and E.E.S. Experiments were performed and analyzed by K.L.P, T.J.C., H.Z., A.B., A.M., C. S., E.S., E.L., A.C-S., M.W., M.B., L.J, and J.L. Animal work was carried out by K.L.P, H.Z., T.J.C., B.F.D, I.S., C.E.S., K.S-C, and A.B. ELISA/Immunohistochemistry was carried out by K.L.P, M.W., M.B., T.J.C., H.Z., D.Q., and B.F.D. HPLC was performed by J.L. Fibrils were obtained from A.W. and K.C.L. E. E. S., C.F.B., K.L.P, T.J.C, D.G. A.W. and K.C.L. gave conceptual advice. T.A.C. and H.B.K. wrote the manuscript.

## Data and materials availability

No large scale data sets were generated in this study. All ASO sequences and chemistries, as well as references to the synthesis are included in the methods to allow for generation of these compounds. SNCA-PAC mice were licensed from Mayo Foundation for Medical Education and Research.

## Acknowledgements

We thank Donna Sipe, Johnnatan Tamayo and Gemma Ebeling for vivarium assistance. Tracy Reigle for figure design and assembly. We also would like to thank the oligo screening group (RTS), the preclinical development team, the oligo synthesis group, vivarium staff and histology core staff at Ionis Pharmaceuticals. We would also like to acknowledge the translational pathology laboratory at Biogen for staining IHC/ISH for the NHP study. We want to acknowledge Mian Horvath, a wonderful technician in the Luk laboratory at UPenn.

## Abbreviations

PD: Parkinson’s disease
CNS: central nervous system
mRNA: mature ribonucleic acid
*Snca/SNCA*: alpha synuclein rodent/human gene, respectively
aSyn: alpha synuclein protein
LB: Lewy body
LN: Lewy neurite
ASO: antisense oligonucleotide
PFF: pre-formed fibril
CSF: cerebrospinal fluid
NHP: non-human primate
MSA: multiple systems atrophy
DLBD: Diffuse Lewy body disease
GD: Gaucher disease
PAF: pure autonomic failure
TH+: tyrosine hydroxylase positive
DA: dopamine
SN: substantia nigra
IHC: immunohistochemistry
pSer129+ aggregates: phospho serine 129 positive aggregates

## Supplementary Material Titles

Fig. S1. ASO1 and ASO2 administration prior to PFF injection result in mRNA reduction and pSer129+ aggregate reduction in multiple rat studies.

Fig. S2. *Snca* mRNA reduction and contralateral TH positive cell counts, from single dose rat ASO administration prior to PFF injection study.

Fig S3. Human *SNCA* ASOs exhibit concentration dependent reduction and a long duration of action.

Fig. S4. Representative images for IHC (anti-ASO and aSyn antibodies) or ISH (*SNCA* mRNA) for motor cortex from NHP (cynomolgus) experiment.

