## Supplemental Files for "Alpha-synuclein antisense oligonucleotides as a disease-modifying therapy for Parkinson’s disease"

### Supplementary Materials

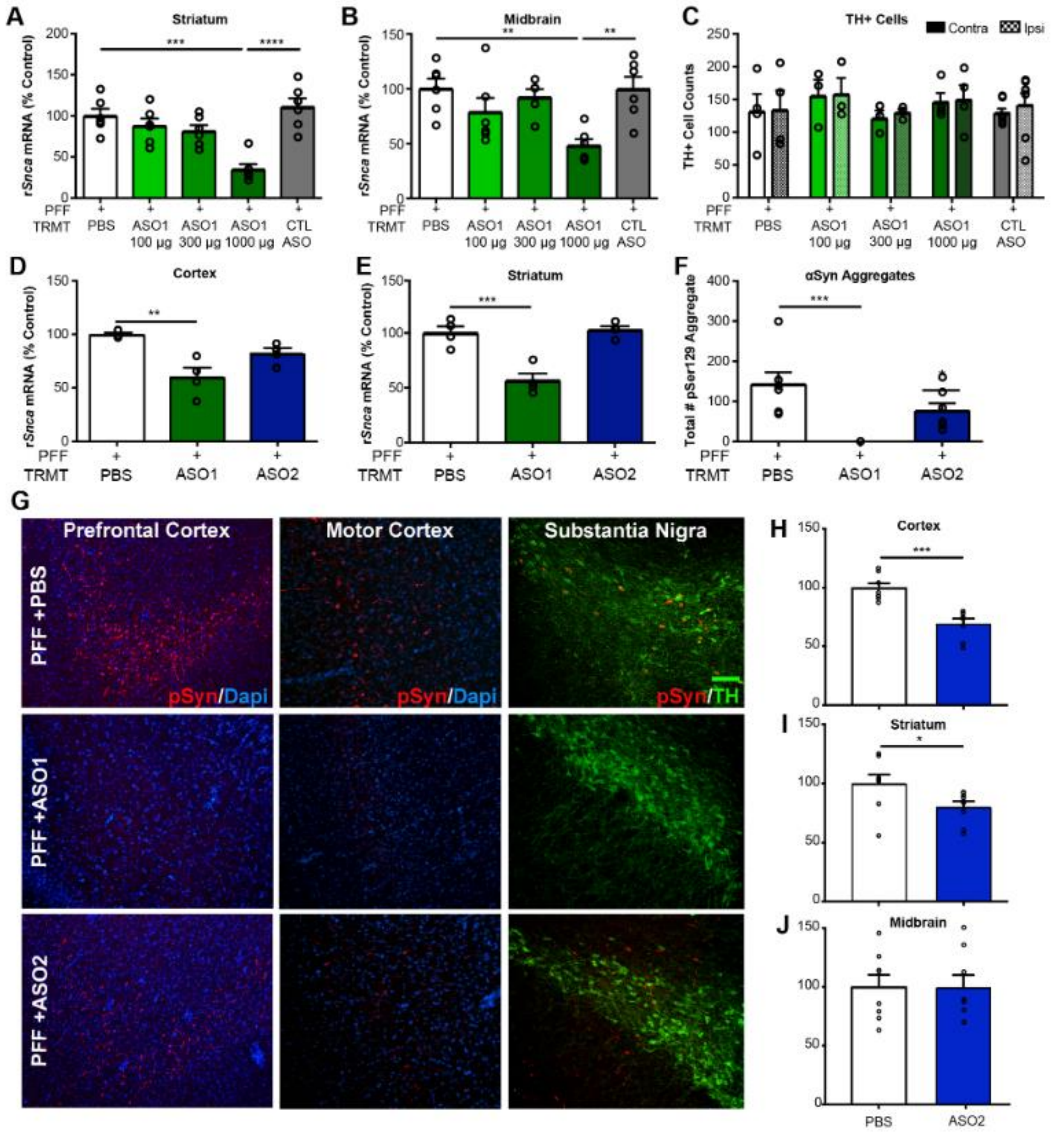


**Fig. S1. ASO1 and ASO2 administration prior to PFF injection result in mRNA reduction and pSyn+ aggregate reduction in multiple rat studies.** (**A** and **B**) mRNA reduction in the striatum and midbrain by RT-PCR for a dose response study in the rat with ASO1 with ASO administration prior to PFF injection with termination at Day 61 (n=6 for all groups) (**C**) Dopaminergic cell counts quantified by motor neuron cell counts (MTN counts) for a dose response study in the rat with ASO1 with ASO administration prior to PFF injection with termination at Day 61. (n=4, 3, 3, 4, 7 for PBS, 100 µg, 300 µg, 100 µg, CTL ASO, respectively) (**D** and **E**) mRNA reduction in the cortex and striatum by RT-PCR from rat PFF study with ASO administration (1000µg) prior to PFF injection comparing ASO1 and ASO2 with termination at day 61 (n=4). (**F**) Quantification of pSyn+ aggregates in the substantia nigra (total enumeration) by IHC from rat PFF study with ASO administration prior to PFF injection comparing ASO1 and ASO2 with termination at day 61 (n=7). (**G**) Representative images of pSyn+ aggregates by immunofluorescence showing prefrontal cortex, motor cortex, and substantia nigra from rat PFF study with ASO administration prior to PFF injection comparing ASO1 and ASO2 with termination at day 61. Scale bar is fluorescent green and equals 100 µm. (**H** to **J**) mRNA reduction in the cortex, striatum, and midbrain by RT-PCR 42 days post a single 1000µg ICV administration of ASO2 (n=4). Data are ± s.e.m. *P<0.05, **P<0.001, ***P<0.0001, ****P<0.00001 (one-way ANOVA with Tukey post hoc analyses). PFF(pre-formed fibril), TRMT(treatment), CTL ASO(Control ASO).

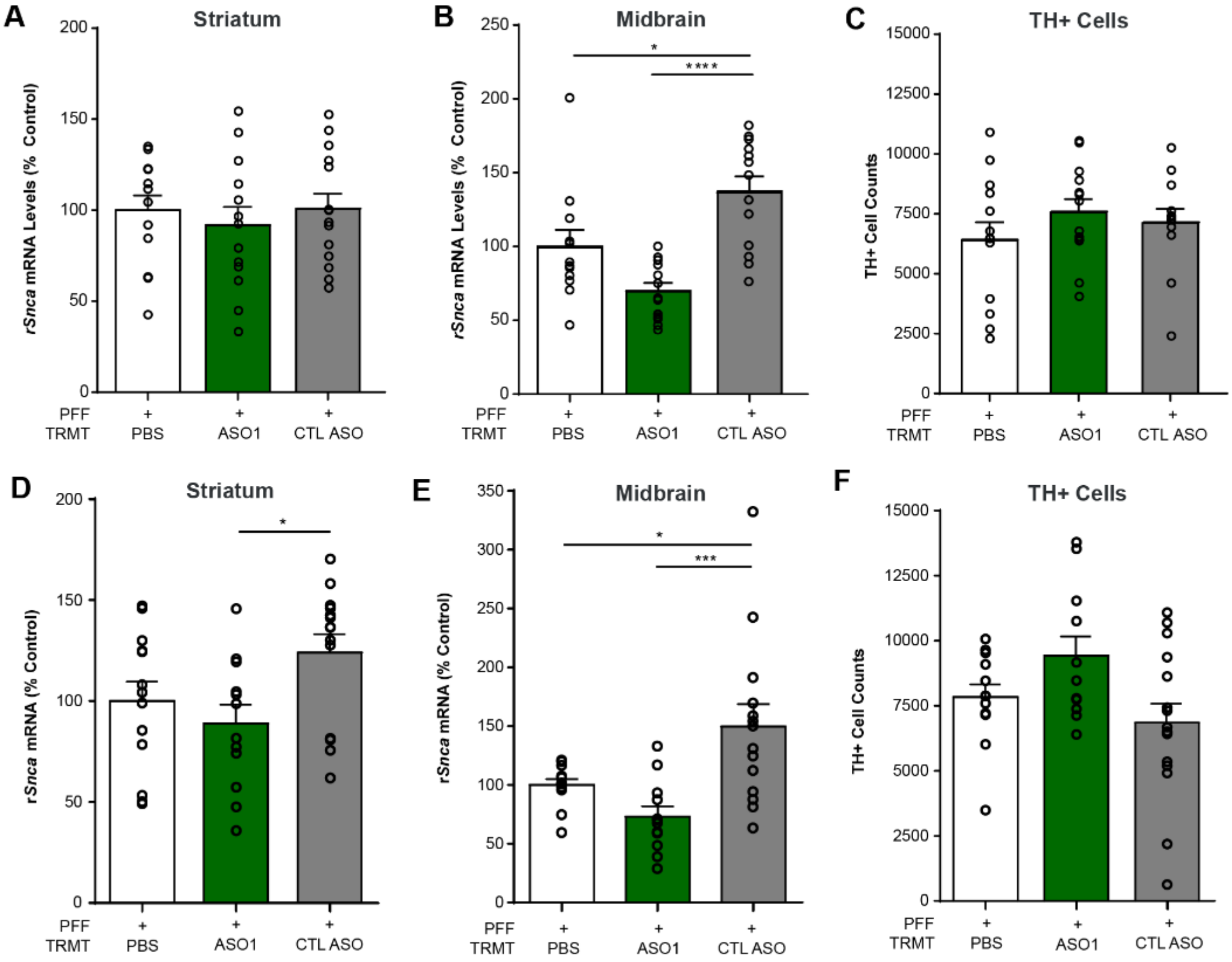


**Fig. S2. *Snca* mRNA reduction and contralateral TH cell counts, from single dose rat ASO administration prior to and after PFF injection.** mRNA in the (**A**) striatum and (**B**) midbrain by RT-PCR for ASO administration prior to PFF injection (n= 13, 13, 14 for PBS, ASO 1, and CTL ASO, respectively) at 181 days post ICV administration. (**C**) Quantification of dopaminergic cell counts in contralateral SN (by stereology) (n=13, 13, 12 for PBS, ASO1, and CTL ASO, respectively). mRNA in the (**D**) striatum and (**E**) midbrain by RT-PCR for ASO administration after PFF injection (n= 13, 12, 14 for PBS, ASO1, and CTL ASO) at 160 days following ASO administration. (**F**) Quantification of dopaminergic cell counts in contralateral SN (by stereology) (n=12, 11, 14 for PBS, ASO1, and CTL ASO, respectively). Data are ± s.e.m. *P<0.05, **P<0.001, ***P<0.0001, ****P<0.00001 (one-way ANOVA with Tukey post hoc analyses).

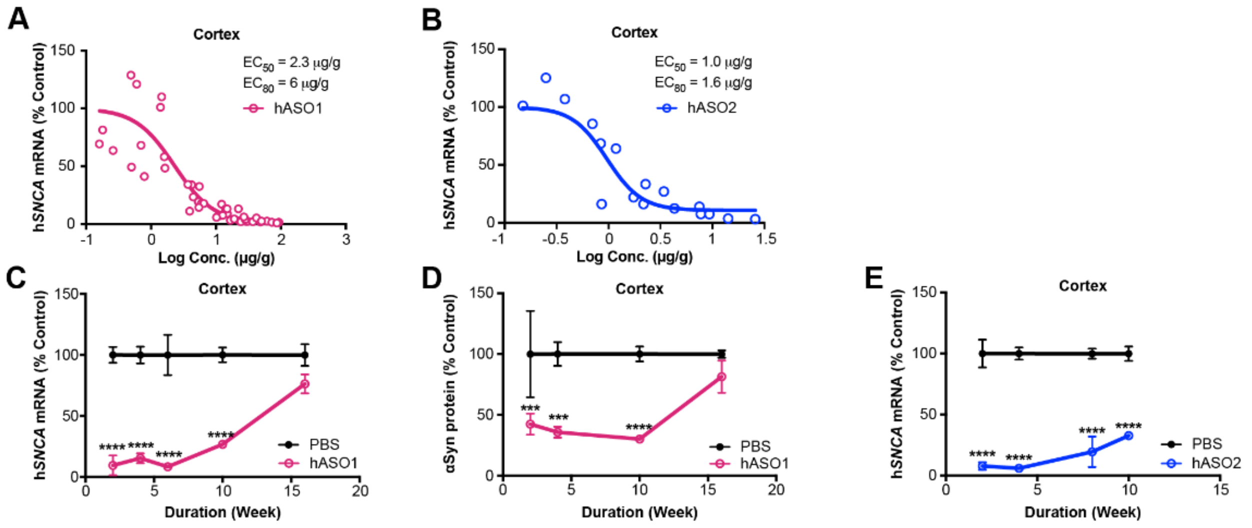


**Fig. S3. Human *SNCA* ASOs exhibit concentration dependent reduction and a long duration of action.** (**A** and **B**) Concentration dependent reduction in *SNCA* mRNA in SNCA-PAC human *SNCA* transgenic mice (n=10 for all groups except 700 µg n=7 for hASO1) (n=8 for all groups for hASO2 (2 mice removed with ROUT analysis from 300 µg and 700 µg groups, combination of two studies of n=4 each, n=12 for all PBS groups). (**C** to **E**) Duration of action of hASO1 and hASO2 (n=4 for hASO1 and hASO2, n= 8, 8, 8, 7, 8 for 2, 3,4, 6, 10, 16 week analysis for PBS administered mice in hASO1 study, n=7, 6, 8, 10 for 2, 4, 8, and 10 week analysis of PBS administered mice in the hASO2 study) for (**C**) mRNA by RT-qPCR for hASO1, (**D**) protein by ELISA for hASO1, and (**E**) mRNA by RT-qPCR for hASO2. Data are ± s.e.m. *P<0.05, **P<0.001, ***P<0.0001, ****P<0.00001 (one-way ANOVA with Tukey post hoc analyses). (Two-way ANOVA with Tukey post hoc analyses for duration of action, repeated measures ANOVA for body weight over time, with all other analyses using One-way ANOVA with Tukey post hoc analyses).

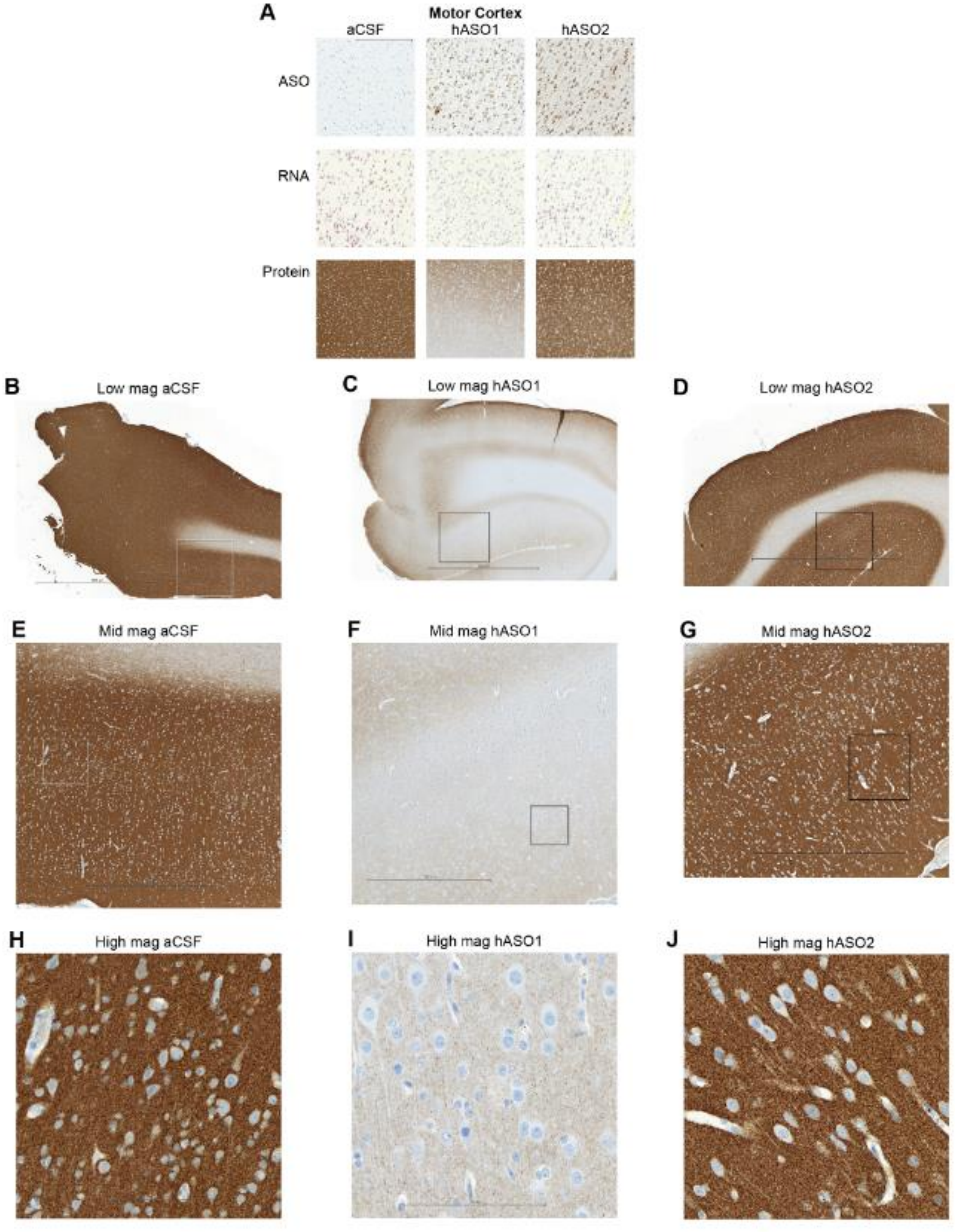


**Fig. S4. Representative images for IHC (anti-ASO and aSyn antibodies) or ISH (*SNCA* mRNA) for motor cortex from NHP (cynomolgus) experiment.** (n=2-4). (**A**) Representative images for IHC (ASO and aSyn antibody) or ISH (*SNCA* mRNA). Scale bar = 600 µm for protein/300 µm for ASO and ISH results for the motor cortex. (**B**) Low magnification (0.625x) of aSyn protein in representative aCSF administered NHP in motor cortex. Inset box indicates magnified image in E. Scale bar = 4500µm. (**C**) Low magnification (0.625x) of aSyn protein in representative hASO1 administered NHP in motor cortex. Inset box indicates magnified image in G. Scale bar = 4500µm (**D**) Low magnification (0.625x) of aSyn protein in representative hASO2 administered NHP in motor cortex. Scale bar = 4500µm. (**E**) Mid magnification (2.5x) of aSyn protein in representative aCSF administered NHP in motor cortex. Inset box indicates magnified image in F. Scale bar = 1000µm (**F**) Mid magnification (2.5x) of aSyn protein in representative hASO1 administered NHP in motor cortex. Inset box indicates magnified image in F. Scale bar = 1000µm (**G**) Mid magnification (2.5x) of aSyn protein in representative hASO2 administered NHP in motor cortex. Inset box indicates magnified image in F. Scale bar = 1000µm (**H**) High magnification (20x) of aSyn protein in representative aCSF administered NHP in motor cortex. Scale bar = 160µm. (**I**) High magnification (20x) of aSyn protein in representative hASO1 administered NHP in motor cortex. Scale bar = 160µm. (**J**) High magnification (20x) of aSyn protein in representative hASO2 administered NHP in motor cortex. Scale bar = 160µm.
